# Defining the optimal intranasal administration strategy for inactivated poliovirus vaccine

**DOI:** 10.1101/2022.07.28.501816

**Authors:** Heleen Kraan, Geert-Jan Willems, Peter C. Soema

## Abstract

Mucosal immunity plays a pivotal role in protection against infection and the transmission of pathogens from person to person. Ideally, novel vaccines should be easy and safe to administer, provide mucosal immunity, be safe to manufacture, and be affordable for low-income countries. Alternative delivery strategies, such as the intranasal route, may fulfill at least some of these preferred vaccine characteristics. Moreover, vaccination via mucosal routes has the potential to evoke strong mucosal immunity at the entry site.

In the current study, the potential of the intranasal route was investigated for an inactivated polio vaccine based on Sabin strains (sIPV). Different vaccination regimes for intranasal administration of sIPV were evaluated by measuring both systemic and mucosal immune responses in mice. Heterologous prime-boost schedules using a combination of parenteral and mucosal administration showed to elicit virus neutralizing antibody titers in serum and polio-specific IgA titers at different mucosal sites in mice that were vaccinated with sIPV. Moreover, the inclusion of an adjuvant was able to further enhance immune responses. Intranasal administration of adjuvanted sIPV combined with a regular intramuscular sIPV vaccination can significantly improve mucosal immune responses while maintaining systemic immune responses, which can lead to better protection against polio infection and possibly prevent virus transmission. An intranasally delivered inactivated polio vaccine might be of value for use in routine vaccination or outbreak control, and therefore, a helpful tool towards polio-eradication.

## 1 Introduction

Mucosal immunity plays a large role in the prevention of infection and subsequent transmission of various infectious diseases, such as polio. While the inactivated polio vaccine (IPV) is effective in disease prevention through the induction of polio-specific systemic antibodies, it does not prevent transmission of the poliovirus. The live attenuated oral polio vaccine (OPV) however, is able to induce polio-specific mucosal immunity, which is a powerful mechanism of protection and interruption of polio transmission. While OPV is thus in theory more effective due to the induction of important mucosal responses against polio, it is currently being phased out by the Global Polio Eradication Initiative due to risk of vaccine-derived polio viruses. Alternative delivery routes for IPV might be able to induce these important mucosal responses [1]. Indeed, mice studies demonstrated that IPV based on Sabin strains (sIPV) administered via the intranasal route was able to induce strong systemic immunity as well as polio-specific IgA antibody responses at multiple mucosal sites, including the intestines, whereas intramuscular immunization with sIPV was unable to do so [2]. Furthermore, the addition of a strong mucosal adjuvant showed to significantly enhance the immune responses elicited upon intranasal sIPV administration [2].

In order to improve the immune responses of IPV administered via the intranasal route, one should not only look at the inclusion of adjuvants, but also consider the dosing regimens of the vaccine. Indeed, heterologous prime-boost approaches with several antigens have shown that the combination of a parenteral and mucosal administration can significantly increase systemic and mucosal immune responses [7–11]. Aside from heterologous prime-boost strategies, fractional dosing has also been used to increase vaccine immunogenicity. Numerous studies have shown that intradermal vaccination with repeated fractional doses increases the immunogenicity of IPV compared to a single full dose of IPV [5]. A study in mice showed that repeated fractional dosing schedules of four times one-fourth (1/4) dose or eight times one-eight (1/8) dose given on consecutive days via the intradermal route elicited significant improved polio-specific IgG titers when compared to those obtained upon vaccination with the same dose given as a bolus either given via the same route (intradermal) or via intramuscular injection [6]. It is possible that the same holds true for intranasal administration.

The main objective of the current study was to determine the optimal vaccination regime for intranasal administration of IPV, which was based on Sabin strains, to induce both potent systemic and mucosal immunity. Multiple heterologous prime-boost combinations as well as fractional dosing were screened using trivalent sIPV in mice. Both functional systemic immunity (serum) as well as polio-specific mucosal IgA antibody responses and B cell responses were determined after vaccination with aforementioned combinations.

## 2 Materials and methods

### 2.1 Materials

Monovalent Sabin IPV (sIPV) bulk material used in this study was produced by Intravacc with an optimized downstream process as described previously [3]. The sIPV bulks used in the dose optimization study were produced at production scale by Sinovac Biotech Ltd. For the preparation of trivalent sIPV, monovalent type 1, type 2 and type 3 were mixed and subsequently concentrated using 10 kDa Amicon® Ultra Centrifugal Filters (Merck Millipore, Billerica, MA) to a nominal concentration of 10000-16000-32000 DU/mL.

The excipients D-Sorbitol and L-glutamic acid monosodium salt monohydrate (MSG) were purchased from Sigma Aldrich (St. Louis, MO), and magnesium chloride hexahydrate (MgCl_2_.6H_2_O) was purchased from Merck (Darmstadt, Germany). Class B CpG oligonucleotide 1826 and polyinosinic-polycytidylic acid (Poly(I:C) HMW) were purchased from InvivoGen.

### 2.2 Immunization studies

Animal experiments were performed according to the guidelines provided by the Dutch Animal Protection Act and were approved by the Committee of Animal Experimentation (DEC) of the National Institute of Public Health and Environment (RIVM). Balb/cOlaHsd mice (6-8 weeks old from Envigo, The Netherlands) received their vaccinations while anesthetized with isoflurane/O_2_.

In general, mice received three times (on day 0, day 7 and day 28) a single human dose (trivalent sIPV: 10-16-32 D-antigen/dose), either via the intramuscular (IM, injection of 50 μL in hind limb) or intranasal (IN, pipetting 5 μL in each nostril) route. The adjuvant dose (CpG or Poly(I:C)) used was 20 μg per dose. Blood samples were taken at day 0 (prior to immunization), day 7, day 14 and day 28. At day 35, mice were sacrificed and relevant samples and organs were isolated.

At the day of sacrifice, anesthetized animals received an intraperitoneal injection of 0.1 mL of 1.8 mM pilocarpine (Sigma-Aldrich, St. Louis, MO) in PBS to induce saliva production. Saliva was collected and, subsequently, animals were sacrificed by bleeding. Post-mortem fecal samples were isolated from the large intestine, weighted and stored at −80°C until analysis. Spleens were placed in RPMI-1640 medium (Gibco, Invitrogen) supplemented with 5% (v/v) fetal bovine serum and placed on ice for B-cell ELISPOT analysis. Small intestines were harvested and placed in 3 mL PBS containing 50 mM EDTA (Gibco, Invitrogen) and protease inhibitors (Complete, Mini, EDTA free, Roche Applied Sciences). Small intestines were extensively vortexed and centrifuged for 15 min at 300 g (4°C). Supernatants, mentioned further as intestinal wash, were collected and stored at −80°C until analysis.

### 2.3 Polio-specific IgG and IgA ELISA

Enzyme linked immunosorbent assays (ELISA) were performed to determine polio-specific antibody titers in sera and intestinal washes. The ELISA was performed as described previously with some small adaptations [2]. In brief, polystyrene 96 wells microtiter plates (Greiner Bio-One, Alphen a/d Rijn, The Netherlands) were coated with bovine anti-poliovirus serum (Bilthoven Biologicals, Bilthoven, The Netherlands) in PBS (Gibco from Invitrogen, Paisley, UK). Plates were washed and trivalent inactivated polio vaccine diluted in assay buffer (PBS supplemented with 0.5% (w/v) skim milk and 0.05% (v/v) Tween 80) was added and incubated for 2h at 37°C. Subsequently, plates were washed and threefold sample dilutions in assay buffer were added and incubated for another 2h at 37°C. After washing, plates were incubated with horse-radish peroxidase (HRP)-conjugated goat-anti-mouse IgG or HRP-conjugated goat-anti-mouse IgA (Southern Biotech, Birmingham, AL). After 1h incubation at 37°C, plates were washed and SureBlue TMB substrate was added. After 10-15 minutes, the reaction was stopped with 0.2M sulfuric acid (Fluka) and absorbance was measured at 450 nm by using a Biotek L808 plate reader. Endpoint titers were determined by 4-parameter analysis using the Gen5™ 2.0 Data Analysis software (BioTek Instruments, Inc., Winooski, VT) and defined as the reciprocal of the serum dilution producing a signal identical to that of negative control samples at the same dilution plus three times the standard deviation.

### 2.4 Virus neutralization (VN) assay

Neutralizing antibodies against all three poliovirus types were measured separately by inoculating Vero cells with 100 TCID50 of the wild-type strains (Mahoney, MEF-1 and Saukett) as described previously [1]. Twofold serial serum dilutions were made and serum/virus mixtures were incubated for three hours at 36°C and 5% CO_2_ followed by overnight incubation at 5°C. Subsequently, Vero cells were added and after 7 days of incubation at 36°C and 5% CO_2_, the plates were stained and fixed with crystal violet and results were read macroscopically. Virus neutralizing (VN) titers were expressed as the last serum dilution that has an intact monolayer (no signs of cytopathogenic effect).

### 2.5 B cell ELISPOT

MultiScreen-HTS IP 96 wells filter plates (Merck Millipore, Darmstadt, Germany) were placed in 35% ethanol, immediately washed with PBS and, subsequently coated overnight with 5 μg/mL monovalent IPV type 1, 2 or 3. As a positive control, wells were coated with a mixture of 7 μg/mL purified goat-anti-mouse kappa and 7 μg/mL purified goat-anti-mouse lambda (Southern Biotech). As a negative control, wells were left uncoated (PBS). After washing with PBS, plates were blocked with RPMI-1640 medium (Gibco, Invitrogen) with 2% (w/v) Protifar (Nutricia, Zoetermeer, The Netherlands) for 1 hour at room temperature. Spleens were homogenized using a 70-μm cell strainer (BD Falcon, BD Biosciences) and cells were collected in RPMI-1640 supplemented with 10% fetal bovine serum (HyClone) and antibiotics (Penicillin-Streptomycin-L-Glutamine, 100x (Gibco, Invitrogen)). Erythrocytes were removed by ACK lysis buffer (Gibco, Invitrogen). After washing, cells were counted and 5×105 cells/well were added to coated plates. After overnight incubation at 37°C and 5% CO2 plates were washed extensively and incubated with HRP-conjugated goat-anti-mouse IgA or goat-anti-mouse IgG (Southern Biotech). Subsequently, BCIP-NBT liquid substrate (Sigma Aldrich, St. Louis, MO) was added and plates were kept in dark during spot development. Thereafter, the reaction was stopped by discarding the substrate and extensively washing of both sides of the filter with tap water. Plates were dried overnight at 37°C and spots were counted using an EliSpot reader.

### 2.6 Statistical analysis

Data was statistically analyzed by comparing all groups by a one-way ANOVA followed by a Tukey’s multiple comparisons test. Probability (*p*) values of *p* <0.05 were considered statistically significant. Statistics were performed using GraphPad Prism version 7.01 (GraphPad Software Inc., La Jolla, CA).

## 3 Results

### Heterologous prime-boost schedules for intranasal sIPV delivery

In order to assess the effect of heterologous prime-boost vaccination on the immune responses elicited by sIPV, mice were immunized according to the schedule depicted in Table 1. As controls, two groups of mice received either an IM or an IN only vaccination with sIPV. Other groups received either an IM prime followed by two IN boosts, or an IN prime followed by two IM boosts. Two different adjuvants, CpG and Poly(I:C), were tested as well with the aforementioned prime-boost combinations.

**Table 1.** Immunization schedule for the study testing heterologous prime-boost immunization schedules combining the intranasal (IN) and intramuscular (IM) routes for sIPV vaccination (study 2).

| Group | n | 1<sup>st</sup> vaccination (day 0)<br>(route – vaccine) | 2<sup>nd</sup> vaccination<br>(route – vaccine) | 3<sup>rd</sup> vaccination<br>(route – vaccine) |
| --- | --- | --- | --- | --- |
| 1 | 10 | IM – placebo | IM – placebo | IM – placebo |
| 2 | 10 | IN – sIPV | IN – sIPV | IN – sIPV |
| 3 | 10 | IM – sIPV | IN – sIPV | IN – sIPV |
| 4 | 10 | IN – sIPV | IM – sIPV | IM – sIPV |
| 5 | 10 | IN – sIPV + CpG | IM – sIPV | IM – sIPV |
| 6 | 10 | IM – sIPV + CpG | IN – sIPV | IN – sIPV |
| 7 | 10 | IN – sIPV + Poly(I:C) | IM – sIPV | IM – sIPV |
| 8 | 10 | IM – sIPV + Poly(I:C) | IN – sIPV | IN – sIPV |

IM – intramuscular; IN – intranasal; CpG – CpG oligodeoxynucleotides; Poly(I:C) – Polyinosinic-polycytidylic acid

Polio-specific IgG titers were measured in serum to determine systemic immune responses. Mice that received a homologous intranasal prime-boost vaccination showed poor to moderate polio-specific IgG titers for all three serotypes (**Figure 1A**). All heterologous prime-boost groups showed significantly higher type 1-specific IgG titers, whereas for type 2 and 3 only mice that received IN prime (followed by IM vaccinations) showed significantly enhanced IgG antibody titers when compared to the IN prime-boost control group. For all serotypes, a significant adjuvant effect of Poly(I:C) was observed upon IM prime vaccination (followed by IN vaccinations) when compared with the group that received IM prime without adjuvant. Only for type 2, an adjuvant effect of CpG was revealed after mucosal prime immunization, not for parenteral prime immunization (**Figure 1A**). As observed in previous studies, polio-specific IgG antibody titers showed to be quite predictable for virus-neutralizing titers as similar results were obtained when assessing functionality of the antibodies by a virus-neutralization assay (**Figure 1B**). Also poor VN titers were observed for the IN prime-boost group with high numbers of non-responders for type 1 and type 2. For type 3, all mice showed functional antibodies after IN prime-boost vaccination. In contrast to the IM prime groups, all IN prime groups, whether or not given with an adjuvant, showed significantly enhanced VN titers against serotype 1 and 2. For type 3, sera from all vaccinated animals showed virus neutralizing capacity with comparable titers for all groups, except the group that received IM prime adjuvanted with CpG, which showed significant higher VN titers when compared to the homologous IN prime-boost (**Figure 1B**). Unexpectedly, serum of some animals in the placebo-group showed to have some virus-neutralizing capacity against type 3. It is unknown what the cause is of these unexpected results (e.g. false positives).

**Figure 1.**
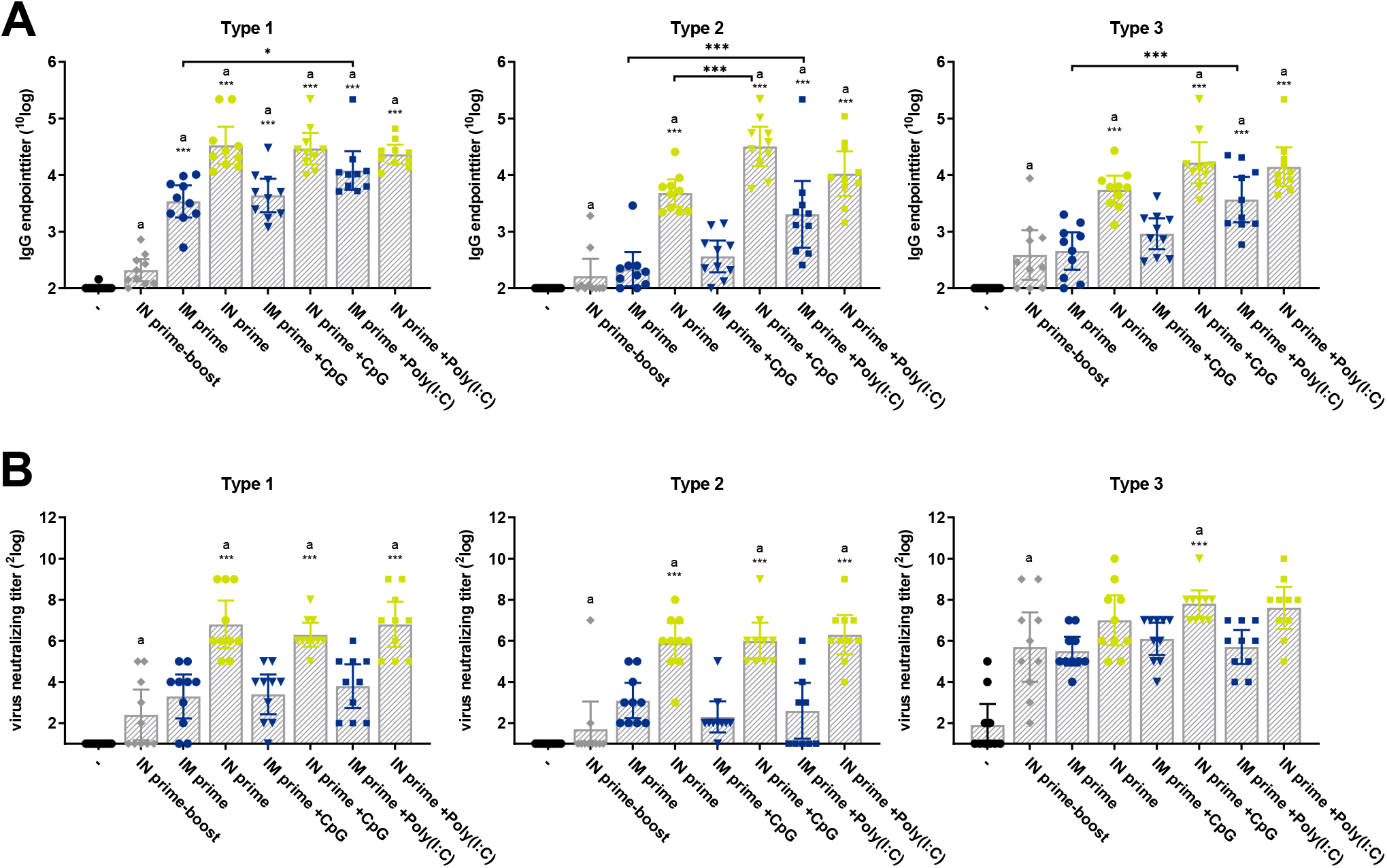
Systemic immunity induced after vaccination with sIPV via heterologous prime-boost vaccination schedules. Polio-specific IgG antibody titers (**panel A**) and virus-neutralizing capacity (**panel B**) of sera from mice (n=10) that received three sIPV vaccinations either via the intranasal route (IN prime-boost) or via heterologous prime-boost vaccination combining intranasal (IN) and intramuscular (IM) routes were measured one week after the third immunization. When ‘IM prime’ is indicated on the x-axis, animals received one IM immunization followed by two IN immunizations. When ‘IN prime’ is indicated, animals received one IN immunization followed by two IM immunizations. Bars depict mean IgG titers or virus neutralizing titers, and error bars show 95% confidence interval values. Asterisks indicate relevant differences between groups (* p<0.05, ** p<0.01, *** p<0.001).

To assess mucosal immunity, polio-specific IgA responses were measured in salivary samples collected one week after the third immunization (**Figure 2**). The groups that received prime immunization via the IM route and booster vaccination via the IN route showed significantly enhanced type 1- and type 3-specific IgA responses compared to the group that received an IN prime and IM boost. The inclusion of Poly(I:C) as adjuvant showed to be beneficial for the induction of type 2- (p<0.05) and type 3-specific IgA antibodies (p<0.001), whereas the inclusion of CpG as adjuvant did not show to be beneficial for the induction of mucosal immunity after heterologous prime-boost with IM prime (**Figure 2**). When starting with an IN prime immunization followed by two IM immunizations, no significantly enhanced polio-specific IgA responses were measured in salivary samples when compared to the placebo-group or homologous IN prime-boost immunization.

**Figure 2.**
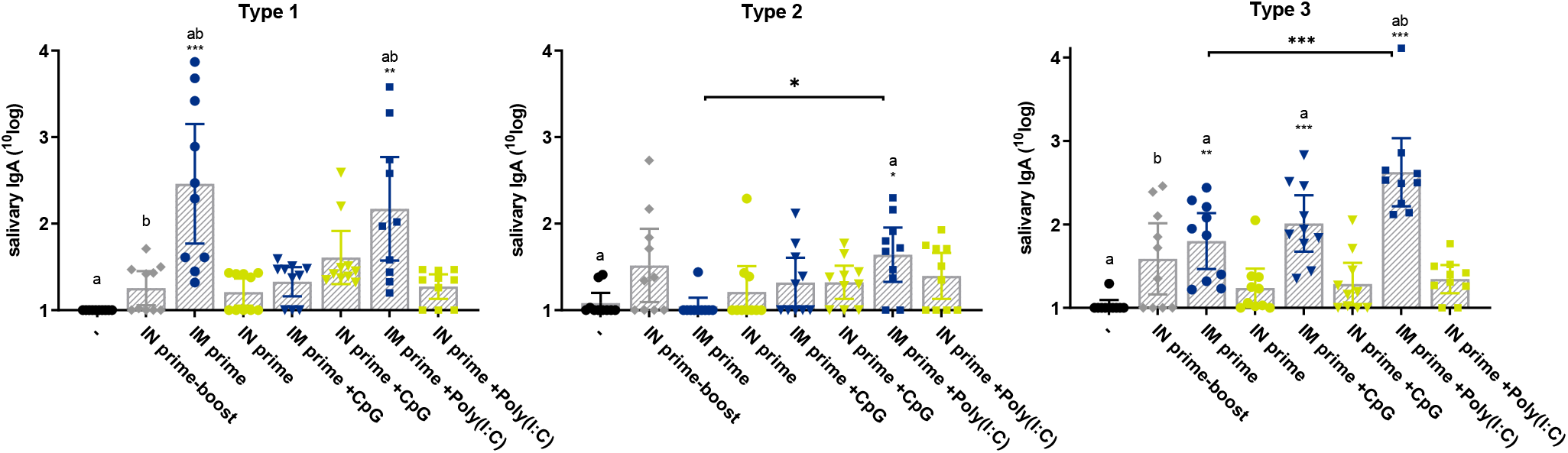
Mucosal immunity induced after vaccination with sIPV via heterologous prime-boost vaccination schedules. Polio-specific IgA antibody titers in salivary samples from mice (n=10), which received three sIPV vaccinations either via the intranasal route (IN prime-boost) or via heterologous prime-boost vaccination combining intranasal (IN) and intramuscular (IM) routes, were measured one weeks after the third immunization. When ‘IM prime’ is indicated on the x-axis, animals received one IM immunization followed by two IN immunizations. When ‘IN prime’ is indicated, animals received one IN immunization followed by two IM immunizations. Bars depict mean IgA titers and error bars show 95% confidence interval values. Asterisks indicate relevant differences between groups (* p<0.05, ** p<0.01, *** p<0.001).

To have a further look at gut immunity induced upon heterologous prime-boost schedules for IN sIPV delivery, polio-specific IgA responses were measured in fecal samples (i.e., extracts from fecal pellets) and intestinal washes, which were collected one week after the third immunization. For the group that received intranasal prime-boost vaccination, all except one type 1-specific IgA antibody titers were below detection limit, whereas all animals (except one for type 2) showed mucosal IgA antibody responses against type 2 and type 3 in fecal samples (**Figure 3A**). Heterologous prime-boost vaccination with sIPV (n the presence or absence of CpG as adjuvant) via IM prime followed by two IN immunizations showed to significantly enhance type 1-specific mucosal immunity in the intestine as measured by fecal IgA antibodies. For type 3, a significant adjuvant effect was observed for Poly(I:C) when given during prime immunization via the IN route (**Figure 3A**). In general, similar observations were made when measuring polio-specific IgA antibodies in intestinal washes. However, the background signal of these samples was higher as depicted by positive IgA titers for the placebo-group (**Figure 3B**). In some cases, a small adjuvant effect was observed for Poly(I:C) when given via IN prime (type 2) or via IM prime (type 3) in a heterologous prime-boost vaccination scheme (**Figure 3B**).

**Figure 3.**
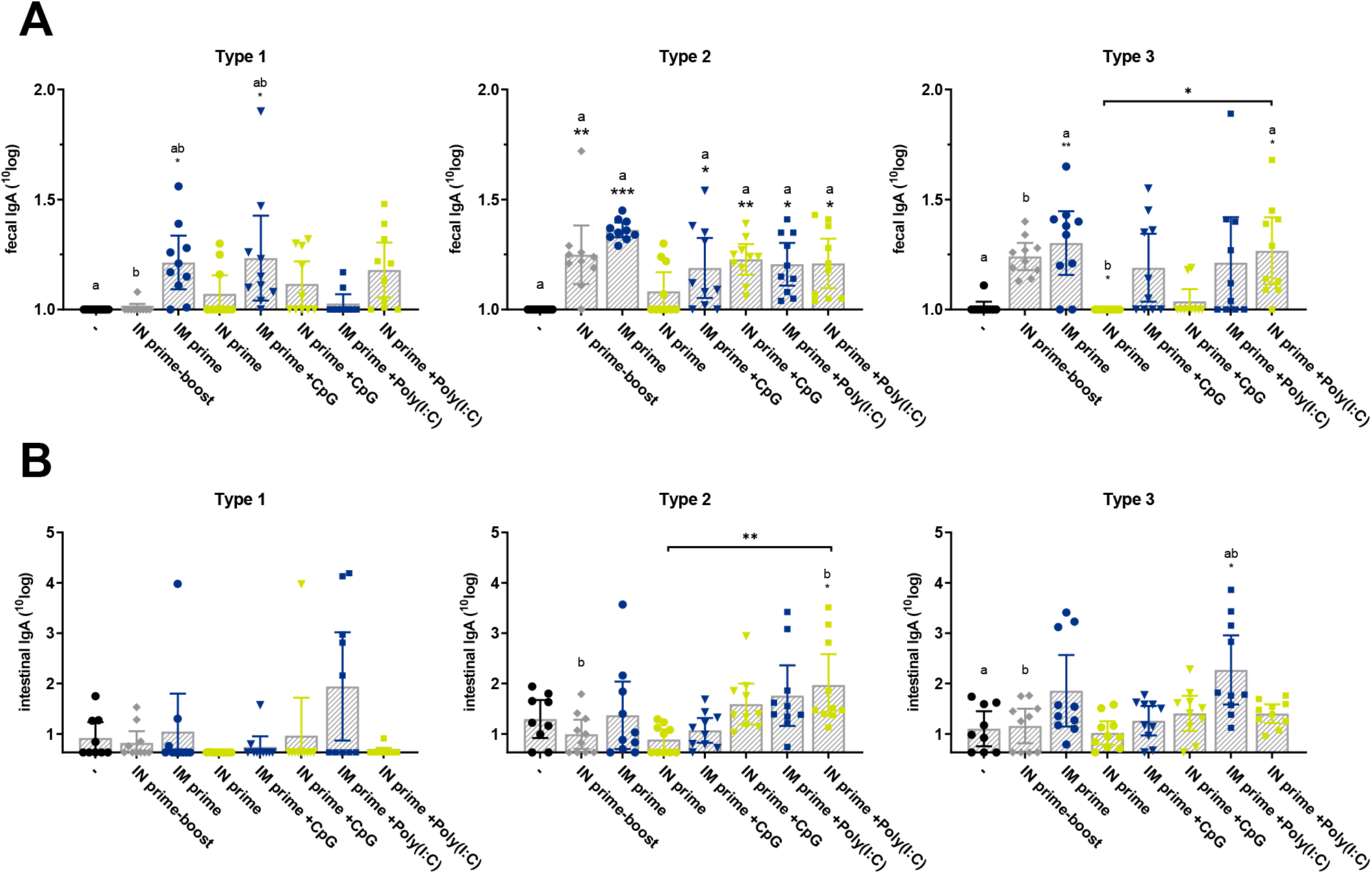
Gut immunity induced after vaccination with sIPV via heterologous prime-boost vaccination schedules. Polio-specific IgA antibody titers in fecal samples (**panel A**) and intestinal washes (**panel B**) from mice (n=10), which received three sIPV vaccinations either via the intranasal route (IN prime-boost) or via heterologous prime-boost vaccination combining intranasal (IN) and intramuscular (IM) routes, were measured one weeks after the third immunization. When ‘IM prime’ is indicated on the x-axis, animals received one IM immunization followed by two IN immunizations. When ‘IN prime’ is indicated, animals received one IN immunization followed by two IM immunizations. Bars depict mean IgA titers and error bars show 95% confidence interval values. Asterisks indicate relevant differences between groups (* p<0.05, ** p<0.01, *** p<0.001).

The effect of homologous versus heterologous prime-boost immunization schedules on the number of polio-specific plasma cells was evaluated in single cell suspensions from spleens. Both the numbers of IgG-producing and IgA-producing B cells were assessed by ELISPOT assay. Irrespective of serotype, all adjuvant-groups showed significant higher numbers of IgG-producing B cells when compared to the placebo-group, but for type 1 these numbers were also higher than the IN prime-boost group (**Figure 4A**). Thus, the inclusion of an adjuvant seemed to stimulate B cells to produce IgG antibodies and, thereby, no clear differences were observed between IN prime versus IM prime for these adjuvant-groups. sIPV supplemented with CpG given either via IN or IM prime showed to significantly enhance the number of type 1-specific IgG-producing B cells when compared to the groups that received sIPV without adjuvant (p<0.01) (**Figure 4A**). The addition of Poly(I:C) as adjuvant in heterologous prime-boost schedules elicited significant higher numbers of IgG-producing splenocytes when compared to those obtained after sIPV vaccination via IN prime-boost. This was irrespective of administration route for prime immunization for type 1 (p<0.001) and type 3 (p<0.01), but also true for type 2 (p<0.001) after IN prime immunization with sIPV plus Poly(I:C) (**Figure 4A**).

**Figure 4.**
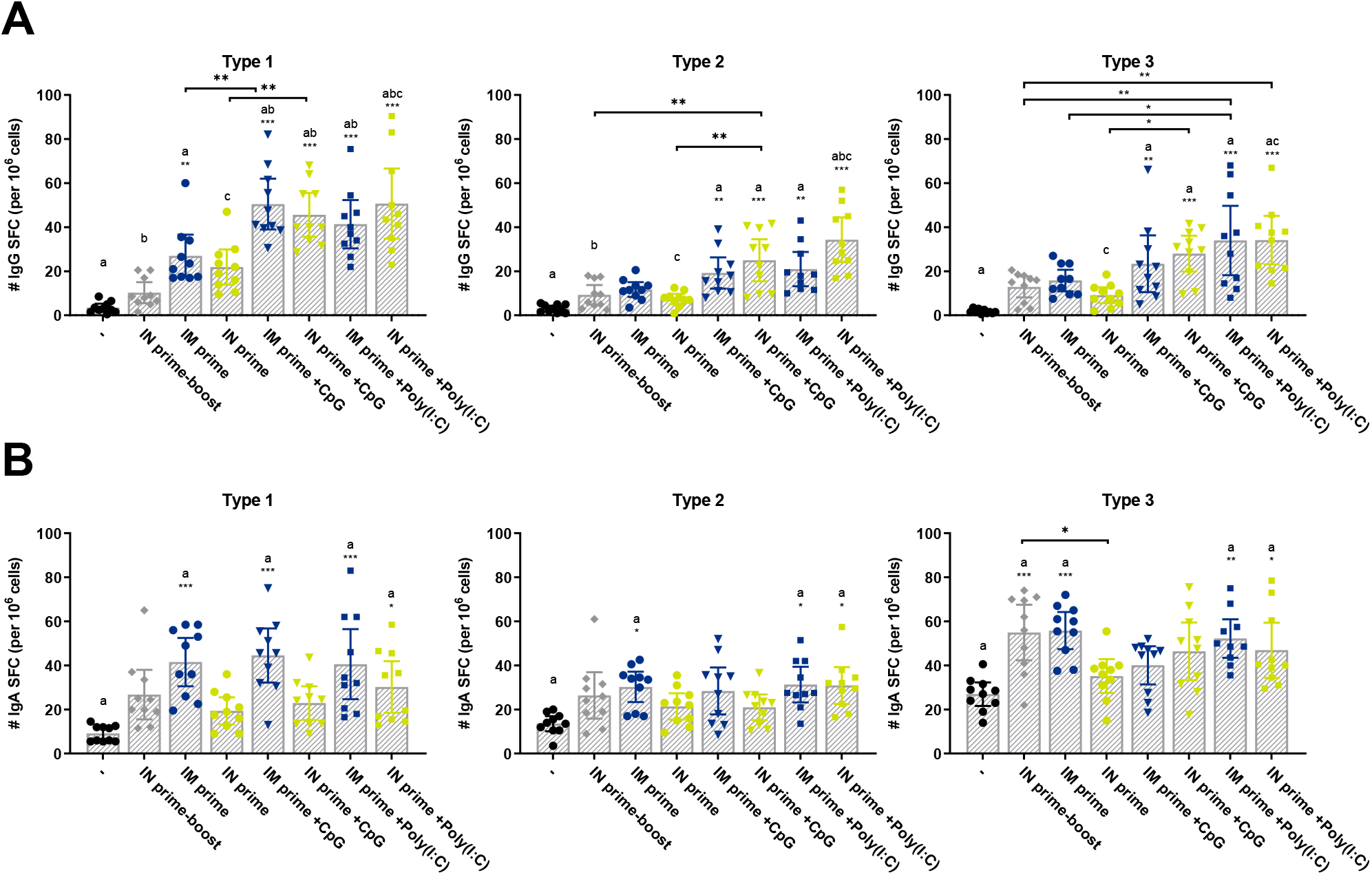
B cell responses elicited after vaccination with sIPV via heterologous prime-boost vaccination schedules. An ELISPOT assay was performed to detect polio-specific IgG-secreting B cells (panel A) and IgA-secreting B cells (panel B) in spleens from mice (n=10) three times immunized with sIPV either via the intranasal route (IN prime-boost) or via heterologous prime-boost vaccination combining intranasal (IN) and intramuscular (IM) routes. When ‘IM prime’ is indicated on the x-axis, animals received one IM immunization followed by two IN immunizations. When ‘IN prime’ is indicated, animals received one IN immunization followed by two IM immunizations. Bars depict mean (antibody-secreting) spot-forming cell (SFC) numbers and error bars show 95% confidence interval values. Asterisks indicate relevant differences between groups (*p<0.05, **p<0.01, ***p<0.001).

As observed for mucosal gut immunity (as measured by polio-specific IgA responses in feces and intestinal washes), highest mucosal immune responses were obtained after heterologous prime-boost immunization starting with an IM prime followed by two immunizations delivered via the IN route (**Figure 3**). ELISPOT data exhibited similar results since highest numbers of IgA-producing B cells were found for groups that received IM prime, whether or not in the presence of an adjuvant (**Figure 4B**). When compared to the number of IgG B cell ELISPOT, the IgA B cell ELISPOT showed a somewhat higher background as illustrated by the numbers of spot-forming cells in the placebo-group. However, still significant higher numbers of IgA-producing cells were observed for IM prime groups (type 1 (p<0,001); type 2 (p<0.05); type 3 (p<0.01)). Also after IN prime with sIPV plus Poly(I:C), increased numbers of IgA-producing cells were observed for the three serotypes (p<0.05) (**Figure 4B**).

### Optimal dosing schedules for intranasal sIPV delivery

The previous experiment showed that heterologous prime-boost immunization schedules combining mucosal and parenteral administration routes can significantly enhance mucosal immune responses. Moreover, studies showed that multiple fractional dosing schedules elicited enhanced immune responses after a single human dose administered via the intramuscular or intranasal route (Supplementary Figures S1-3). Therefore, a follow-up study was designed in which both immunization schedules will be combined resulting in a heterologous prime-boost regime starting with an intramuscular (IM) prime followed by two intranasal immunizations given via multiple fractional doses. Although best results were obtained giving four fractional doses at four subsequent days (4 times ¼ dose), such a schedule would be difficult to implement in the field. For that reason, it was decided to include multiple fractional doses given via the nose on two subsequent days (2 times ½ dose). As the inclusion of an adjuvant showed to be beneficial, Poly(I:C) was included in this study; this adjuvant performed slightly better than CpG in the previous immunogenicity study. An overview of all groups included in this immunogenicity study is shown in **Table 2**.

**Table 2.** Control and test groups included in the study testing different sIPV formulations via heterologous prime-boost immunization schedule using intramuscular (IM) priming and multiple fractional intranasal (IN) doses (study 3).

| Group | n | 1<sup>st</sup> vaccination<br>(route – vaccine – dosing) | 2<sup>nd</sup> vaccination<br>(route – vaccine – dosing) | 3<sup>rd</sup> vaccination<br>(route – vaccine – dosing) |
| --- | --- | --- | --- | --- |
| 1 | 10 | IM – placebo | IM – placebo | IM – placebo |
| 2 | 10 | IM – sIPV – 1 shd | IM – sIPV – 1 shd | IM – sIPV – 1 shd |
| 3 | 10 | IM – sIPV – 1 shd | IN – sIPV – 2x$\frac{1}{2}$ shd | IN – sIPV – 2x$\frac{1}{2}$ shd |
| 4 | 10 | IM – sIPV + Poly(I:C) – 1 shd | IN – sIPV + Poly(I:C) – 2x$\frac{1}{2}$ shd | IN – sIPV + Poly(I:C) – 2x$\frac{1}{2}$ shd |

IM – intramuscular; IN – intranasal; form – formulation; sIPV – Sabin inactivated poliovirus vaccine; shd – single human dose (10-16-32 DU/dose)

To measure systemic immune responses, polio-specific IgG titers were measured in serum. Mice that received conventional homologous IM vaccination showed high polio-specific IgG titers for all three serotypes (**Figure 5A**). All animals that were vaccinated via heterologous prime-boost with multiple fractional intranasal doses induced detectable polio-specific IgG responses against all serotypes. However, these IgG antibody titers were significantly lower than those obtained upon IM vaccination. The inclusion of Poly(I:C) as adjuvant showed to be beneficial for the induction of type 2- and type 3-specific IgG antibodies (**Figure 5A**). As observed in previous studies, polio-specific IgG antibody titers showed to be quite predictable for virus neutralizing titers as similar results were obtained when assessing functionality of the antibodies by a virus neutralization assay (**Figure 5B**).

**Figure 5.**
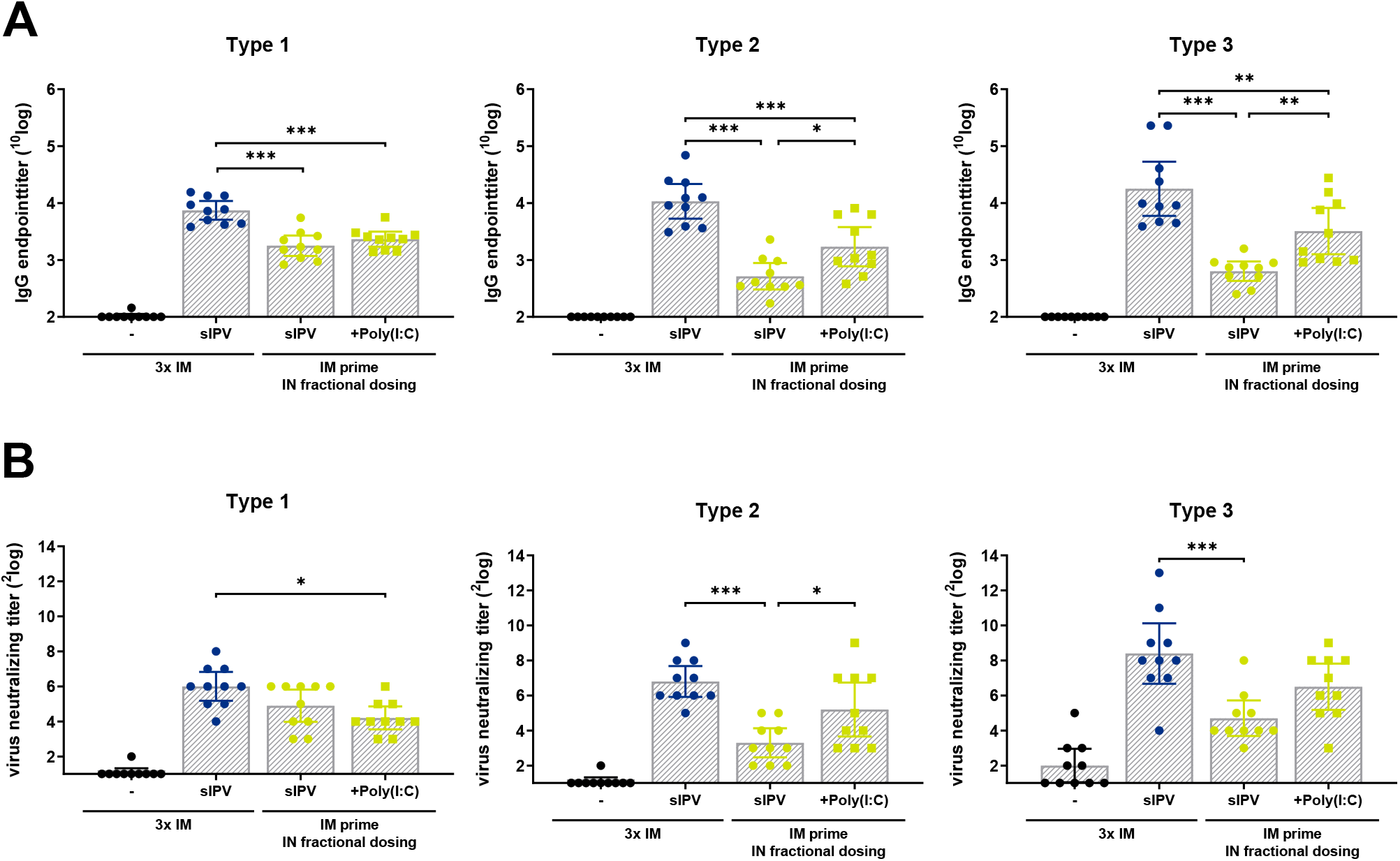
Systemic immunity induced after vaccination with sIPV via heterologous prime-boost immunization using multiple fractional doses given via intranasal route. Polio-specific IgG antibody titers (**panel A**) and virus-neutralizing titers (**panel B**) of sera from mice (n=10) that received three sIPV vaccinations either via the conventional intramuscular (IM) route or via IM prime followed by two intranasal (IN) vaccination given via multiple fractional doses, were measured one week after the third immunization. Bars depict mean IgG titers (panel A) or mean VN titers (panel B), and error bars show 95% confidence interval values. Asterisks indicate relevant differences between groups (* p<0.05, **p<0.01, ***p<0.001).

All vaccinated animals showed detectable virus neutralizing antibody titers and highest titers were obtained upon homologous IM vaccination. For type 1, all heterologous prime-boost groups showed comparable virus neutralizing titers, which were all significantly lower than those observed in the IM group. The inclusion of an adjuvant was not beneficial for the induction of type 1-specific functional antibodies, but a valuable effect of Poly(I:C) on anti-type 2 and anti-type 3 functional antibodies was observed (**Figure 5B**). Unfortunately, serum of some animals in the placebo-group showed to have some virus-neutralizing capacity against type 3. It is unknown what the cause is of these unexpected results (e.g. false positives).

To assess mucosal immunity, polio-specific IgA responses were measured in salivary samples, which are collected one week after the third immunization. As expected, no (type 1 and type 3) to very poor (type 2) mucosal immunity, as measured by polio-specific salivary IgA antibody titers, was induced upon homologous IM vaccination (**Figure 6**). Heterologous prime-boost with fractional IN dosing with the conventional sIPV formulation, whether or not administered in the presence of Poly(I:C), elicited detectable polio-specific IgA antibody titers, which are above background level (all serotypes), but also significantly higher than those obtained after IM vaccination (type 1 and type 3). The inclusion of Poly(I:C) as adjuvant showed to be beneficial for the induction of IgA antibodies against type 2 and type 3 (**Figure 6**). In order to assess gut immunity, polio-specific IgA antibody responses were measured in fecal samples. As expected, highest numbers of responders were observed for the group vaccinated with regular sIPV plus Poly(I:C) as adjuvant via a heterologous prime-boost immunization schedule (**Figure 7**). This sIPV plus Poly(I:C) group showed significantly enhanced type 1- and type 3-specific IgA responses (p<0.01) in fecal samples.

**Figure 6.**
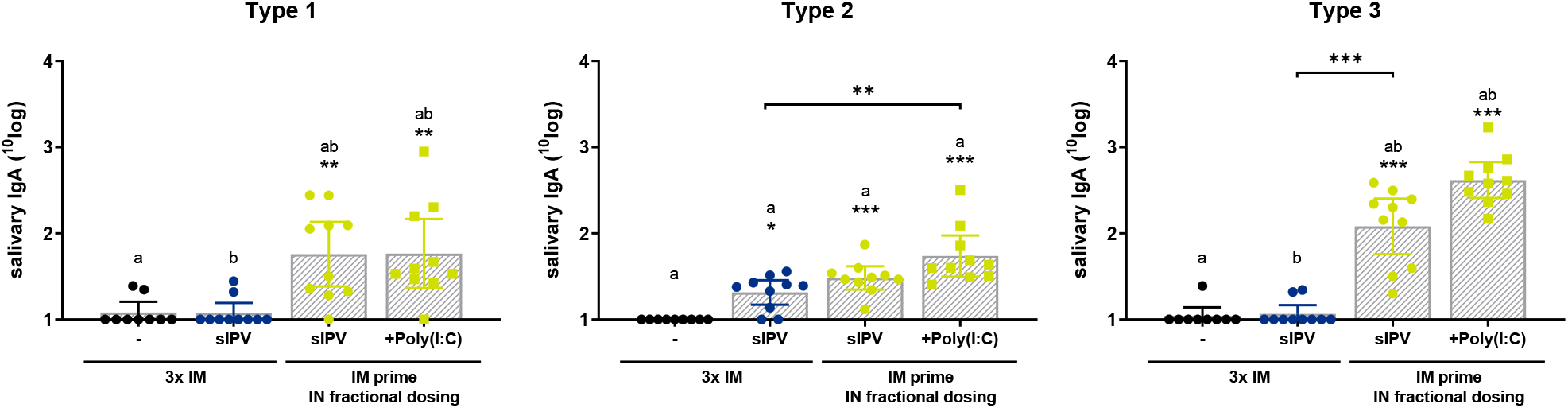
Mucosal immunity induced after vaccination with sIPV via heterologous prime-boost immunization using multiple fractional doses given via intranasal route. Polio-specific IgA antibody titers in salivary samples from mice (n=10) that received three sIPV vaccinations either via the conventional intramuscular (IM) route or via IM prime followed by two intranasal (IN) vaccinations given via multiple fractional doses, were measured one week after the third immunization. Bars depict mean IgA titers and error bars show 95% confidence interval values. Asterisks indicate relevant differences between groups (* p<0.05, ** p<0.01, *** p<0.001).

**Figure 7.**
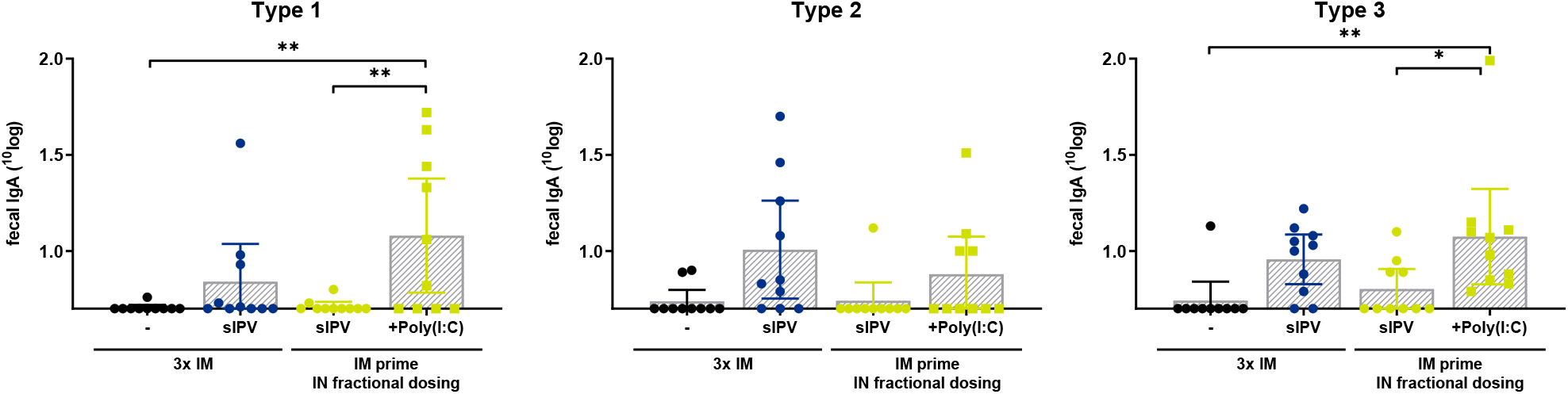
Gut immunity induced after vaccination with sIPV via heterologous prime-boost immunization using multiple fractional doses given via intranasal route. Polio-specific IgA antibody titers in fecal samples from mice (n=10) that received three sIPV vaccinations either via the conventional intramuscular (IM) route or via IM prime followed by two intranasal (IN) vaccinations given via multiple fractional doses, were measured one week after the third immunization. Bars depict mean IgA titers and error bars show 95% confidence interval values. Asterisks indicate relevant differences between groups (*p<0.05, **p<0.01, ***p<0.001).

The effect of homologous conventional IM vaccination versus a heterologous prime-boost with multiple fractional IN dosing schedule on the number of polio-specific plasma cells was evaluated in single cell suspensions from spleens. Both the numbers of IgG-producing and IgA-producing B cells were assessed by ELISPOT assay. Very poor numbers of IgG-producing B cells were observed upon 3xIM vaccination or after IM prime and fractional IN vaccination using sIPV without adjuvant (**Figure 8A**). In the presence of Poly(I:C), significant enhanced numbers of IgG-producing B cells against type 2 (p<0.001 and p<0.01) and type 3 (p<0.05 and p<0.001) were induced. sIPV plus adjuvant showed significant higher numbers of type 2-specific (p<0.05) and type 3-specific (p<0.05) IgG-secreting B cells than those activated after three IM vaccinations or heterologous prime-boost vaccination of plain sIPV (p<0.05 for type 2 and p<0.01 for type 3) (**Figure 8A**).

**Figure 8.**
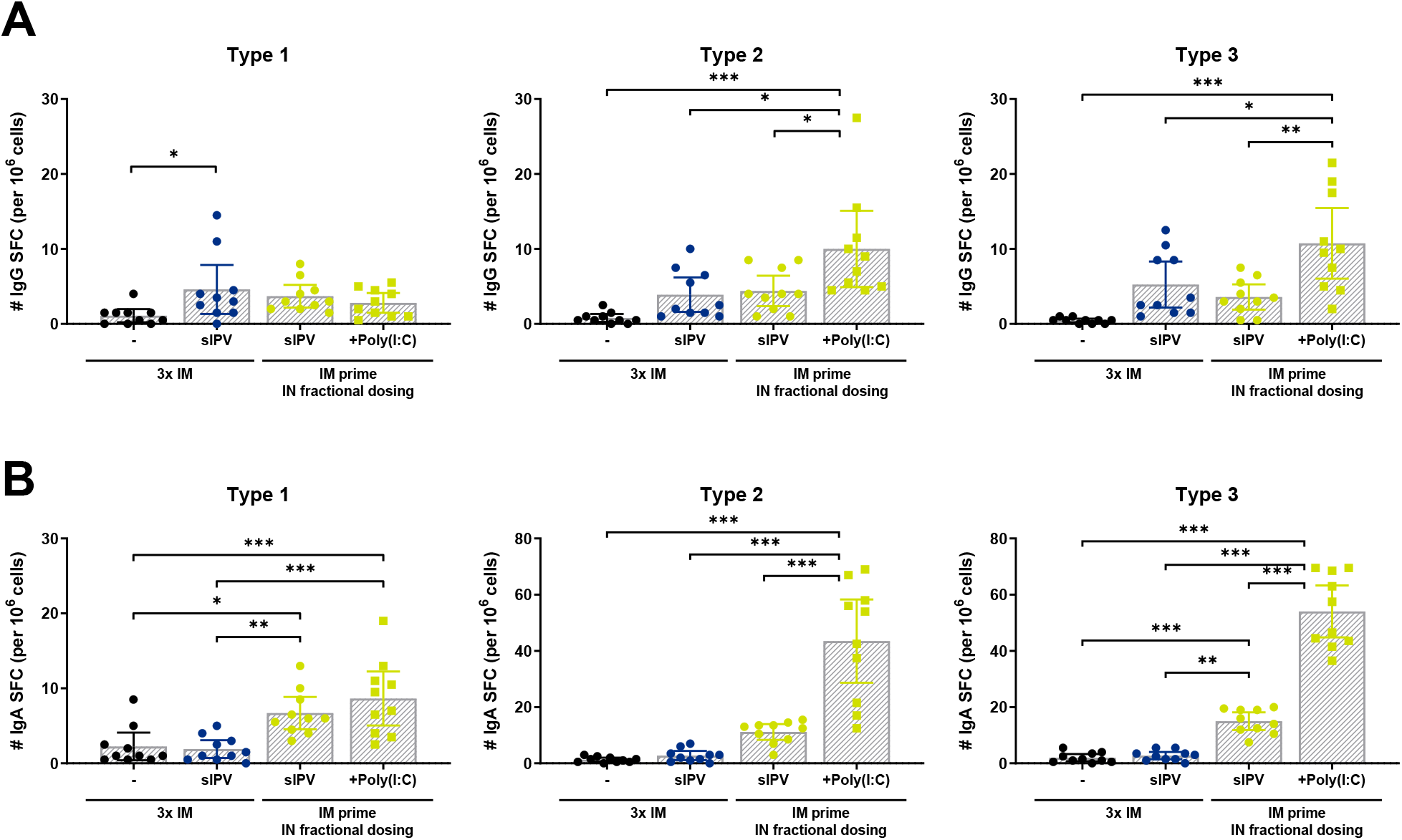
B cell responses elicited after vaccination with sIPV via heterologous prime-boost immunization using multiple fractional doses given via intranasal route. An ELISPOT assay was performed to detect polio-specific IgG-secreting B cells (panel A) and IgA-secreting B cells (panel B) from spleens from mice (n=10) three times immunized with sIPV either via the conventional intramuscular (IM) route or via IM prime followed by two intranasal (IN) vaccinations. Bars depict mean (antibody-secreting) spot-forming cell (SFC) numbers and error bars show 95% confidence interval values. Asterisks indicate differences between groups (*p<0.05, **p<0.01, ***p<0.001).

As observed for mucosal immunity (as measured by polio-specific IgA responses in saliva and fecal samples), highest mucosal immune responses were obtained after heterologous prime-boost with multiple fractional IN dosing of sIPV plus Poly(I:C) as adjuvant (**Figure 6** and **Figure 7**). ELISPOT data exhibited similar results since numbers of IgA-producing B cells for the sIPV plus Poly(I:C) group were significantly above background level for all serotypes (p<0.001), but also significantly higher than the IM group for all serotypes (p<0.001) (**Figure 8B**).

## 4 Discussion and conclusion

Induction of local mucosal immune responses against viruses is key to prevent virus infection and transmission. With OPV being phased out, alternatives based on inactivated polio vaccines should be developed that are able to induce mucosal immunity. Administration of sIPV via the intranasal route has shown promising results in mice previously when combined with adjuvants, and was explored further in the current study.

The main objective of this study was to determine the optimal vaccination regime for intranasal administration of sIPV to induce both systemic and mucosal immunity. Both multiple fractional dosing and heterologous prime-boost immunization regimes were evaluated. In contrast to OPV delivered via the oral route, mucosal polio vaccination based on IPV required the inclusion of an adjuvant to elicit appropriate immunity against polio, which was already confirmed in a previous study in mice [1]. Literature on the use of a mucosal route for IPV is limited; one study previously explored the use of a double mutant of LT (dmLT) in combination with the sublingual route [4]. However, toxin-based adjuvants are not recommended for use via the intranasal route as this might cause unwanted adverse effects (Bell’s palsy) [5, 6]. Several adjuvants that have shown their potential for (Sabin) IPV via the parenteral route could also be evaluated for mucosal vaccination, like LPS-derivate PagL [7], CAF01 [8] or chitosan [9]. However, for a fast track to market, adjuvants with a proven safety record in humans might be preferred over components with immune potentiating capacity or delivery systems that are not licensed yet. From this point of view, CAF01 (phase I), flagellin (phase II), saponins (phase II), CpG oligonucleotides (phase III), or poly(I:C) (phase III), or outer membrane vesicles (phase I) [10, 11] are worthwhile to test in combination with IPV delivered via mucosal routes. In this study, both CpG ODN and Poly(I:C) were evaluated, as these components showed already their adjuvant capacity in combination with sIPV delivered via the mucosal route in earlier studies (unpublished data).

Poly(I:C) induced higher polio-specific systemic IgG, salivary and intestinal IgA responses compared to CpG ODN in the heterologous prime-boost study. In the experiment with heterologous prime-boost combined with fractional dosing, sIPV adjuvanted with poly(I:C) managed to induce superior numbers of IgA-producing B cells compared to the other groups, clearly indicating the benefit of the addition of the adjuvant. Indeed, poly(I:C) has shown its potency as an intranasal adjuvant for influenza vaccines in previous studies with mice and pigs [12, 13]. Since poly(I:C) is already in phase III trials, it is a suitable candidate for an intranasally applied sIPV vaccine.

The initial immunogenicity study clearly showed that heterologous prime-boost schedules have potential for sIPV vaccination. When compared to homologous prime-boost using the intranasal (IN) route, heterologous prime-boost vaccination starting with IN prime showed to significantly enhance the induction of systemic IgG antibodies, which also showed to be able to neutralize the poliovirus. However, a drawback of this immunization schedule was the poor induction of mucosal IgA antibodies in the intestine. In contrast, the heterologous prime-boosting schedule starting with intramuscular (IM) prime followed by two IN immunizations was able to induce similar or even better systemic immunity for all three types and also comparable to improved mucosal immunity. Vaccination via mucosal routes generally needs relatively high antigen doses to circumvent tolerance mechanisms. Therefore, multiple fractional dosing schedules might not the best strategy for intranasal sIPV administration. However, the first study revealed that this vaccination regime might has potential for both the intramuscular and intranasal route by showing increased systemic immune responses after multiple fractional doses when compared to the same dose given at once. Nevertheless, our results showed that a single unadjuvanted dose is not enough to induce potent immune responses upon intranasal vaccination. The inclusion of an adjuvant might be necessary and a prime-boost vaccination regime can also further enhance systemic and mucosal immune responses.

The follow-up study was designed based on the results from the initial experiments. As multiple fractional dosing might improve systemic immunity and heterologous prime-boosting was able to improve both systemic and mucosal immunity when compared to homologous intranasal sIPV vaccination, it was decided to combine both strategies. Conventional homologous IM vaccination was compared with heterologous prime-boost including IM prime and multiple fractional doses given via the IN route. As expected, strong systemic immunity, but undetectable to very poor mucosal immunity was elicited upon IM vaccination. The heterologous prime-boost with fractional dosing schedule was able to significantly enhance mucosal immunity as measured by salivary polio-specific IgA antibody titers. Moreover, slightly lower systemic immunity was induced when compared to IM vaccination, but still strong polio-specific IgG titers with virus neutralizing capacity were elicited. Best results were obtained for the conventional sIPV formulation supplemented with Poly(I:C) as adjuvant. Furthermore, this group showed highest numbers of anti-polio IgG- and IgA-producing B cells.

This study showed that intranasal administration of inactivated polio vaccine combined with a regular intramuscular vaccination can significantly improve mucosal immune responses, which in turn can lead to better protection and, perhaps more importantly, the prevention of further transmission of the virus. Intranasally administered inactivated polio vaccines could therefore be an important addition in the array of polio vaccines that help us eradicate the polio virus.

## Acknowledgments

This work was supported by the World Health Organization as Global Polio Eradication Initiative (GPEI) collaborative research project for the Polio Research Committee.

