## Supplementary data for "Defining the optimal intranasal administration strategy for inactivated poliovirus vaccine"

### Background

Numerous studies have shown that intradermal vaccination with repeated fractional doses increases the immunogenicity of IPV compared to a single full dose of IPV [12]. A study in mice showed that repeated fractional dosing schedules of 4 times ¼ dose or 8 times 1/8 dose given on consecutive days via the intradermal route elicited significant improved polio-specific IgG titers when compared to those obtained upon vaccination with the same dose given as a bolus either given via the same route (intradermal) or via intramuscular injection [13]. It is possible that the same holds true for intranasal administration. Although not tested for the intradermal route, it might be interesting to check whether this effect is also visible when giving one dose divided over two consecutive days (instead of 4 or 8 days) since this might be more practical and therefore easier to implement.

### Experimental design

In the first immunization study, mice received a single human dose (trivalent sIPV: 10-16-32 DU/dose) as bolus or via multiple fractional doses at subsequent days via the intramuscular (IM, injection of 50 µL in hind limb) or intranasal (IN, pipetting 5 µL in each nostril) route. Blood samples were taken at day 0 (prior to immunization), day 7 and day 14. At day 21, mice were sacrificed and relevant samples and organs were isolated (see section **Error! Reference source not found.**).

***Table S1** – Immunization schedule for the study testing multiple fractional doses for sIPV both via the intramuscular (IM) and intranasal (IN) route (study 1).*

| Group | n | Route | Vaccine | Volume | Dose[1] | Vaccination schedule: |
|---|---|---|---|---|---|---|
| **1** | 10 | IM | Placebo | 0.05 mL | - | Day 0 |
| **2** | 10 | IM | sIPV | 0.05 mL | 1 shd | Day 0 |
| **3** | 10 | IM | sIPV | 0.05 mL | ½ shd | Day 0 – Day 1 |
| **4** | 10 | IM | sIPV | 0.05 mL | ¼ shd | Day 0 – Day 1 – Day 2 – Day 3 |
| **5** | 10 | IN | sIPV | 10 µL | 1 shd | Day 0 |
| **6** | 10 | IN | sIPV | 10 µL | ½ shd | Day 0 – Day 1 |
| **7** | 10 | IN | sIPV | 10 µL | ¼ shd | Day 0 – Day 1 – Day 2 – Day 3 |

*[1]Dose of a single immunization. In total all mice received the same dose. 1 single human dose (shd) means 10-16-32 DU.*

### Results: multiple fractional dosing

### Systemic immunity

To measure systemic immune responses, polio-specific IgG titers and virus-neutralizing titers were measured in serum. Mice that received multiple fractional sIPV doses via the intramuscular route, either 2 times ½ dose or 4 times ¼ dose, induced significant improved systemic immune responses when compared with animals that received the same dose given in once as bolus injection (**Figure S1**). Unfortunately, the number of responders and IgG titers in the intranasal groups were low. Only for type 3, a significant improved IgG titer was observed for the group vaccinated four times with ¼ dose when compared with the other intranasal groups (**Figure S1A**). Similar results were obtained by testing functionality of serum antibodies as measured by virus-neutralizing capacity of the serum. No measured type 1 and type 2 virus-neutralizing titers in the groups immunized via the intranasal route, whereas higher numbers of responders were observed when measuring type 3 virus-neutralizing antibody titers in the groups vaccinated via multiple fractional doses (2x ½ shd and 4x ¼ shd) when compared to one single human dose given as bolus via the intranasal route (**Figure S1B**). This indicates that the multiple fractional dosing schedule have potential for the intranasal route. Probably the inclusion of an adjuvant or a booster vaccination regime can further enhance immune responses after intranasal sIPV delivery.

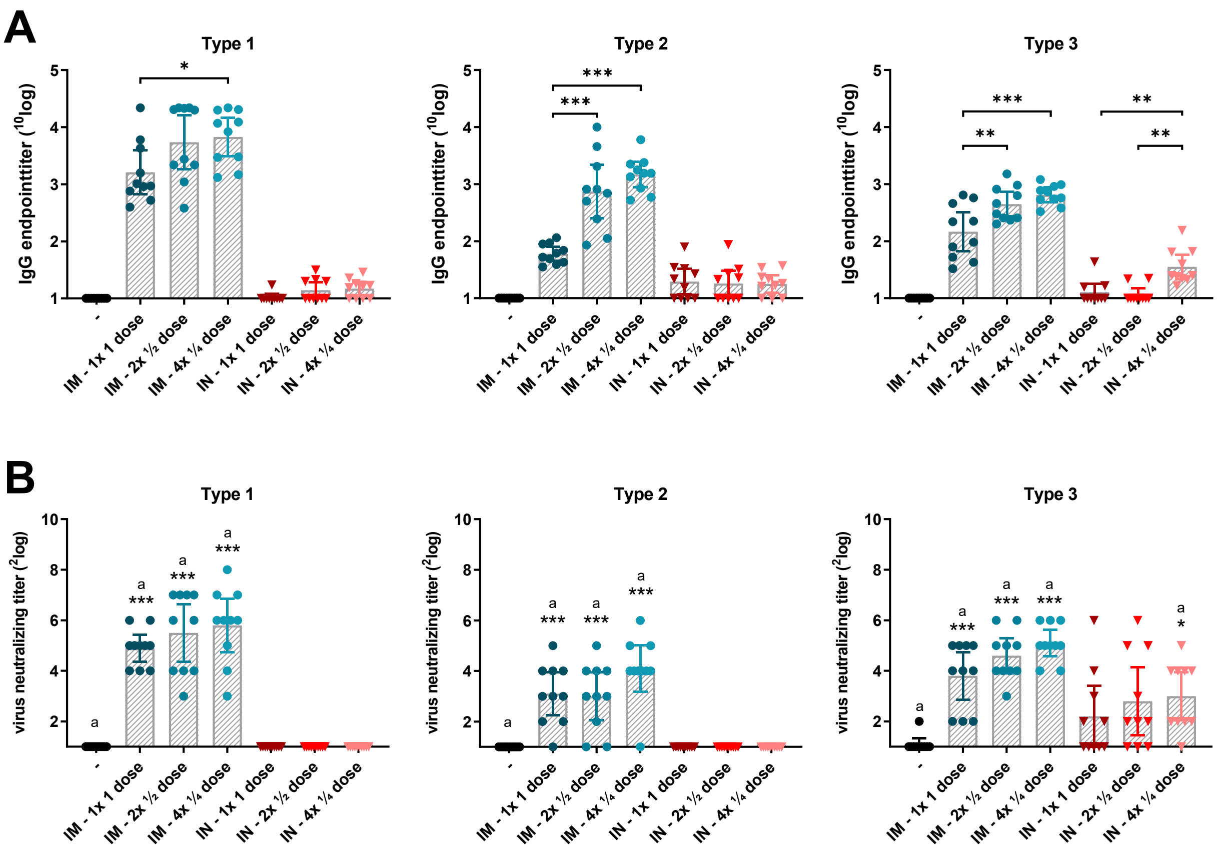


***Figure S1 – Systemic immunity induced after multiple fractional sIPV doses.*** *Polio-specific IgG antibody titers* ***(panel A)*** *and virus neutralizing capacity (****panel B****) of sera from mice (n=10) three weeks after immunization with one single human dose sIPV administered either as bolus (1x 1 dose) or as multiple fractional doses (2x ½ dose; 4x ¼ dose) via conventional intramuscular (IM) injection or via the intranasal (IN) route. Bars depict mean IgG titers or virus neutralizing titers, and error bars show 95% confidence interval values. Asterisks indicate differences between groups (* p<0.05, ** p<0.01, *** p<0.001).*

### Mucosal immunity

To assess mucosal immunity, polio-specific IgA responses were measured in salivary samples (**Figure S2**). As expected based on the poor systemic immune responses observed for the intranasal groups, almost no detectable polio-specific IgA antibody titers can be found. Although some animals showed IgA responses above background level, none of the groups showed significantly enhanced mucosal immune responses when compared with the control group that received placebo vaccine. In addition to the saliva, IgA antibodies were measured in intestinal washes to assess possible gut immunity. From previous work, we know that highest polio-specific IgA responses can be found in salivary samples from mice immunized with sIPV. Thus, as expected all polio-specific IgA responses measured in intestinal washes were at background level (data not shown).

### B cell responses

The effect of multiple fractional dosing on the number of polio-specific plasma cells was evaluated in single cell suspensions from spleens. Both the numbers of IgG-producing and IgA-producing B cells were assessed by ELISPOT assay (**Figure S3**). Significant enhanced numbers of type 1-specific IgG-producing B cells were observed in the intramuscular groups (**Figure S3A**). No significant differences were observed among the intramuscular groups, but a trend was visible that multiple fractional doses resulted in slightly lower numbers of type 1-IgG producing B cells. Unexpectedly, none of the intramuscular groups showed type 2- and type 3-specific IgG-producing B cells (**Figure S3A**). As already revealed from IgA ELISAs on mucosal samples, no enhanced numbers of polio-specific IgA-producing B cells were observed in this study (**Figure S3B**).

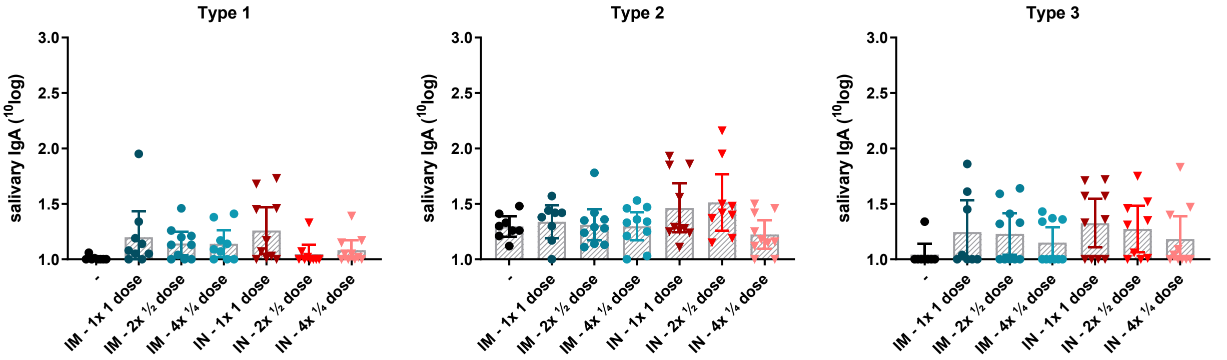


***Figure S2 – Mucosal immunity induced after multiple fractional sIPV doses.*** *Polio-specific IgA antibody titers in salivary samples from mice (n=10) immunized with one single human dose sIPV administered either as bolus (1x 1 dose) or as multiple fractional doses (2x ½ dose; 4x ¼ dose) via conventional intramuscular (IM) injection or via the intranasal (IN) route. Bars depict mean IgA titers and error bars show 95% confidence interval values. No significant differences between groups were observed.*

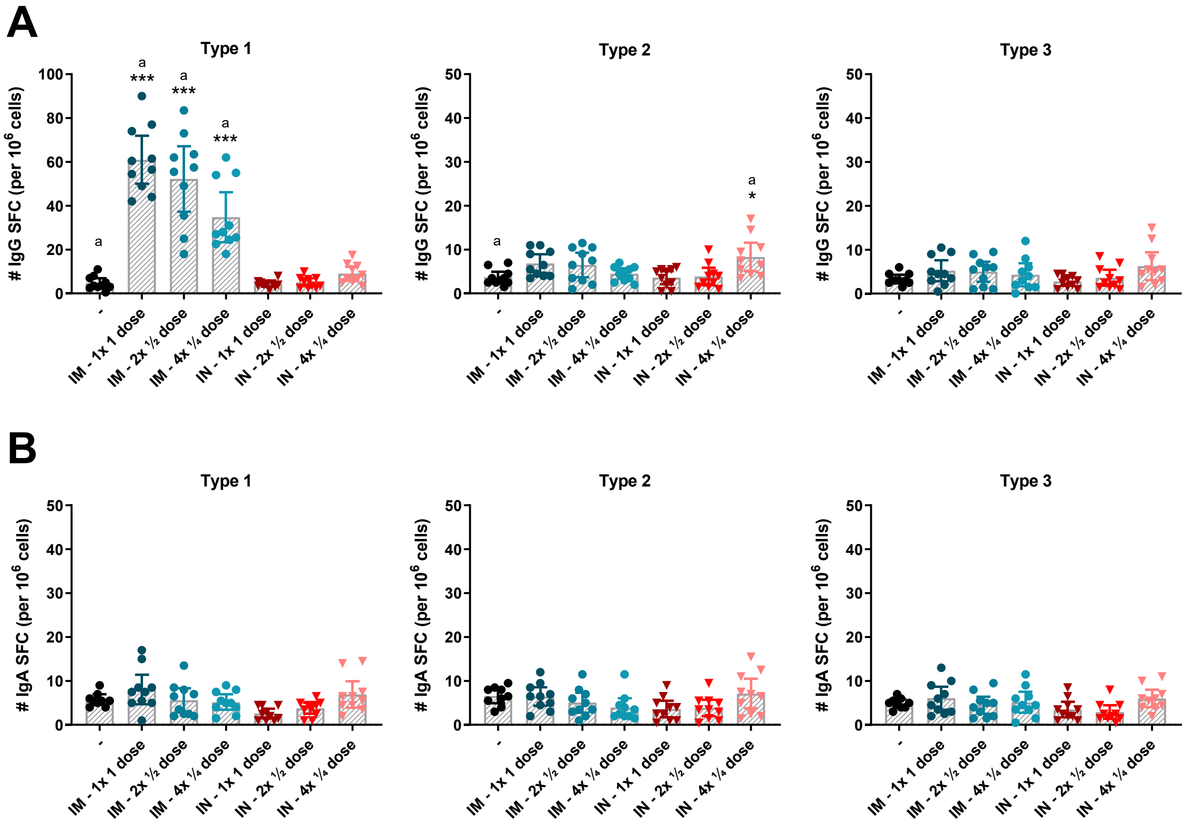


***Figure S3 – B cell responses elicited after multiple fractional sIPV doses.*** *An ELISPOT assay was performed to detect polio-specific IgG-secreting B cells (**panel A**) and IgA-secreting B cells (**panel B**) in spleens from mice (n=10) after immunization with one single human dose sIPV administered either as bolus (1x 1 dose) or as multiple fractional doses (2x ½ dose; 4x ¼ dose) via conventional intramuscular (IM) injection or via the intranasal (IN) route. Bars depict mean (antibody-secreting) spot-forming cell (SFC) numbers and error bars show 95% confidence interval values. Asterisks indicate relevant differences between groups (* p<0.05, ** p<0.01, *** p<0.001).*
